# SCREWx: A Screwless, Chronic, Recoverable, and Lightweight Neuropixels fixture for freely-moving rodents

**DOI:** 10.64898/2026.01.06.697790

**Authors:** Alexandra Cheng, Tate DeWeese, Yue-Qing Zhou, Yotaro Sueoka, Sai Koukuntla, Merrill Green, Kathleen Cullen, James Knierim, Austin Graves, Timothy Harris

## Abstract

High-density Neuropixels probes enable the study of large neural populations with single-cell and sub-millisecond resolution. While single-probe and acute head-fixed experiments have yielded critical scientific insights, understanding the neural mechanisms underlying many complex behaviors requires simultaneous multi-region recordings in freely moving, chronically implanted animals. Various probe fixtures have been developed to enable high-density recording, but existing designs impose critical limitations: their substantial weight restricts the maximum probe count that smaller animals can support, their bulky dimensions constrain the proximity of targeted brain regions, and their complex assembly risks damaging the probe during insertion and recovery. In this paper, we present a lightweight, fully 3D-printable, compact, and screwless fixture for chronic Neuropixels implants in freely moving rodents that features simple mechanisms for stable implantation and safe extraction. Our fixture design enables stable, high-yield single-unit recordings for months-long experiments, along with an 83% successful probe extraction rate. This fixture design provides a robust and accessible solution for long-term, multi-probe chronic Neuropixels recordings, increasing experimental throughput and enabling more complex experimental designs to investigate brain-wide neural dynamics.

## Introduction

Deeper understanding of complex behaviors and cognition requires monitoring large-scale neural dynamics across multiple brain regions in freely-moving animals. Neuropixels probes ^1,2^ (Fig 1), a high-density silicon-based electrophysiology probe, have revolutionized the field by enabling simultaneous recording from 384 channels with single-cell, sub-millisecond resolution^3,4^. While acute experiments using Neuropixels recordings in head-fixed animals have transformed our knowledge on topics such as sensory integration^5,6^ and decision-making^7,8^, utilization of chronic, multi-probe Neuropixels recordings in freely moving behavioral models has remained a challenging feat because of the lack of proper tools supporting stable recordings for extended periods, thus limiting our understanding of brain-wide neural dynamics underlying many naturalistic behaviors. The substantial cost of Neuropixels probes makes it economically unsustainable to perform a single-use implantation for every animal (i.e., cementing probes in place). The need for tools that enable the reuse of these expensive probes across experiments is increasingly apparent as experiments using multiple probes become more common. To address this demand, researchers have developed a variety of implantation strategies to enable probe recovery and reuse^9–13^. Most commonly, these utilize implantable fixtures that encase the probe and are affixed to the animal’s skull, providing a protective envelope from which the probe can be safely extracted after the experiment concludes.

**Figure 1:**
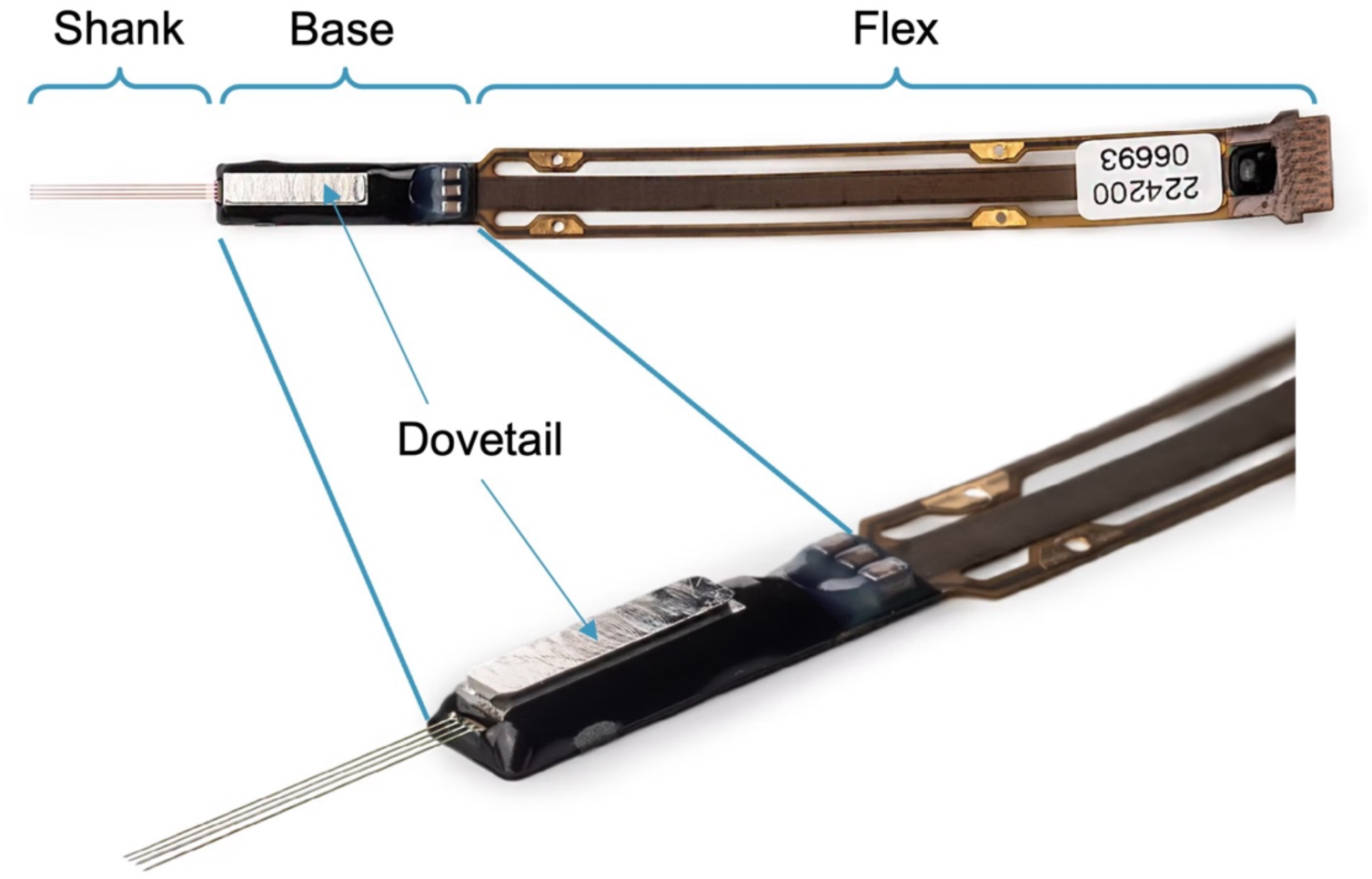
Anatomy of Neuropixels 2.0 multi-shank metal cap probes. The probe consists of three main parts: (1) Shanks: thin multi-pronged feature housing the recording electrode sites that insert into the brain (2) Base: rectangular stiff structure from which the shanks extend and which houses the integrated circuitry underneath the metal dovetail cap (3) Flex: flexible flat cable structure that connects with a printed circuit board for recording and transmission of data

Currently published reusable fixture designs pose challenges that limit their widespread adoption. First, some designs are rather **heavy and bulky**, hindering naturalistic behaviors in smaller animal models such as mice, as well as restricting the number and proximity of multi-probe placements. Second, their **insertion and extraction methods are complex and unreliable**. Many rely on a surgeon’s manual dexterity or complex manipulations (e.g., screws, adhesives)—all of which exert off-axis forces to the probe, posing a significant risk to the fragile probes and surrounding neural tissue.

To overcome these obstacles, we present a **novel, lightweight, and compact fixture** for chronic Neuropixels implants: ***SCREWx***. (Fig 2) **(Github Link: https://github.com/NP-HarrisLab/Harris-Lab-Neuropixels-Fixture-Design)** Our design features a screwless, interlocking three-part assembly that securely holds the probe and isolates it from adhesives applied to the skull, while enabling a low-force extraction along the long axis of the probe, minimizing strain to the fragile shanks. The fixture interfaces with specialized holders for precise, automated control over both implantation and extraction. All fixture elements can be readily made in-house using a lab-grade 3D stereolithography (SLA) printer, making these designs and the experiments they enable accessible to any lab. This novel design directly addresses the critical challenges of weight, size, and handling safety, making complex multi-region chronic recordings from freely moving animals more accessible and scalable.

**Figure 2:**
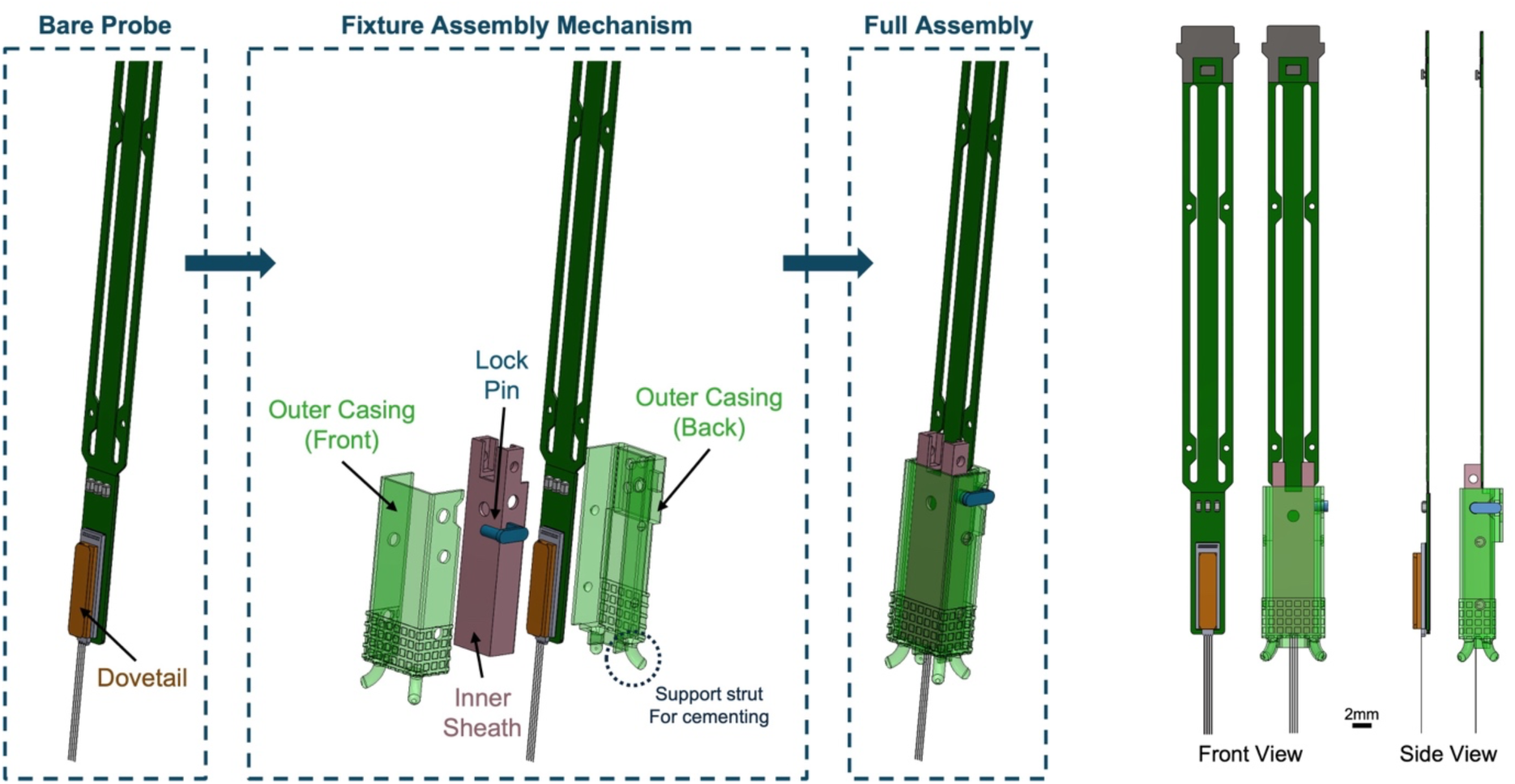
SCREWx design overview. The fully assembled structure measures 6.3mm x 3.6mm x 15.5mm (excluding the optional flaring cement leg structures on the bottom)

## Results

### Fixture design and assembly

The SCREWx fixture design is implemented for Neuropixels 2.0 multi-shank metal cap probes, capitalizing on the probe’s lightweight and compact footprint. While we have analogous designs compatible with Neuropixels 1.0 probes, the 2.0 design represents the optimal implementation and is the focus of this paper. All fixture components are printed using an SLA 3D printer (Form 3B+, Formlabs, Massachusetts, USA). The fixture adds only 0.9 mm of width to each long side and 0.6 mm to each short side of the bare probe, and weighs approximately 0.4 g. The assembly process consists of three straightforward steps, each involving the addition of one of three primary components: (1) an inner sheath to hold the probe via the metal dovetail, (2) a two-part outer casing for skull adhesion, and (3) a removable lock pin to secure the fixture assembly.

***Step 1: Probe mounting in inner sheath***. Assembly starts with sliding the probe into its inner sheath, which features guide rails whose dimensions precisely match those of the probe’s metal dovetail (Fig 3a). The probe is fully seated when its base is flush with the sheath’s edge. Critically, this inner sheath is mechanically isolated from all other components and is the only part recovered post-experiment.

**Figure 3:**
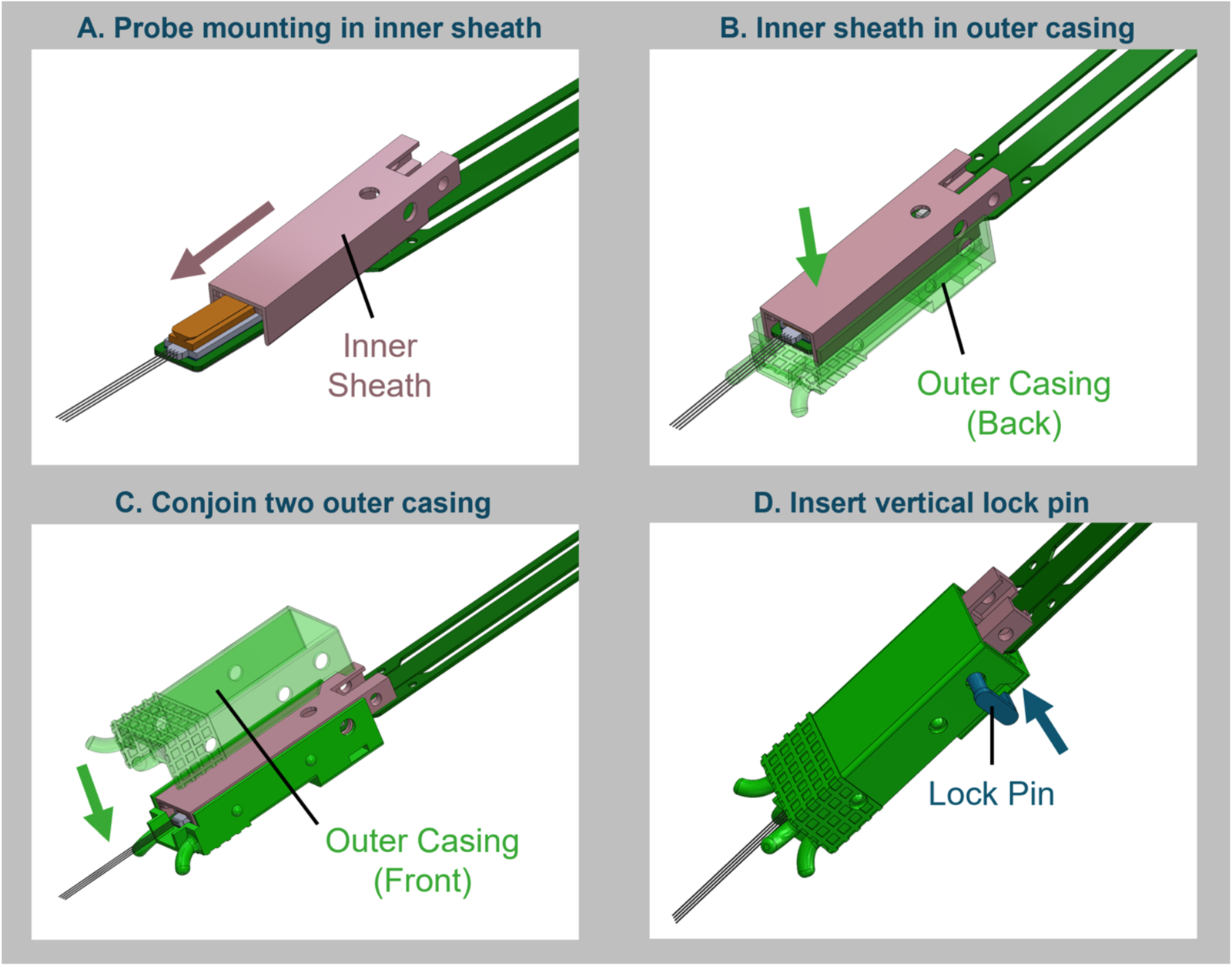
SCREWx assembly process. (a) The metal dovetail (orange) on the probe is slid into the corresponding groove of the inner sheath (pink) (b) The inner sheath-probe assembly is carefully lowered into the bottom half outer casing (translucent green) (c) The top half outer casing (translucent green) is snapped onto the bottom half (solid green), and the four nubs/cutouts on the sidewalls provide precise alignment for mating the two pieces (d) Properly assembled inner sheath and outer casing pieces will reveal an aligned circular cutout that allows for insertion of a small lock pin (blue), restricting relative movement between the probe and all fixture pieces and securing the assembly

***Step 2: Encapsulation with two-part outer casing.*** Next, a two-part outer casing is assembled *around* the inner sheath. This split-shell design avoids sliding components over the delicate shanks, minimizing the risk of damage. The two halves are fastened together via a snap-fit mechanism: small protrusions on one piece lock securely into corresponding holes on the other (Fig 3b & 3c). This mechanism also serves to self-align the two halves, ensuring proper seating and further protecting the shanks from misalignment damage during closure of the pieces. This outer casing provides an adhesion surface to securely anchor the fixture to the skull while fully protecting the mechanically isolated inner sheath-probe assembly. To enhance the adhesion process, the casing’s exterior incorporates customizable support struts whose geometry can be modified to suit specific implant trajectories.

***Step 3: Securing the assembly with the lock pin.*** Finally, a compact 3D-printed lock pin secures the entire assembly, preventing relative motion between the probe, inner sheath, and outer casing. The pin features two key extrusions: a circular shaft that passes through all components to lock them together vertically, and a rectangular key that fits into a corresponding slot on the outer casing to prevent rotation (Fig 3d). This completed assembly will then be mounted on the implantation rod to be inserted into the brain as a whole entity.

### A simple screwless implantation mechanism

Precise implantation of Neuropixels probes requires a micromanipulator to ensure a slow, controlled insertion speed to minimize damage to brain tissue and the probe. This approach necessitates a probe holder to rigidly mount the fixture assembly to the manipulator without angular deviation. Many existing systems rely on screws to secure the fixture to the insertion holder. While secure, this common approach introduces significant risk when releasing the probe-fixture assembly. Once the fixture is cemented to the skull, the implanted probe behaves as a cantilever beam; the torque required to unfasten the screw translates directly into shear and torsional stress on this fixed beam, applying forces along the most fragile axis of the probe shanks, risking probe damage along with damage to the surrounding brain tissue.

To eliminate this critical failure point, we designed a screwless “tongue-and-groove” mounting system. The system consists of a simple, 3D-printable probe holder with a keyed tongue at its distal end (Fig 4a). This part of the probe holder slides securely into the corresponding groove integrated into one of the outer casings of the fixture. When fully seated, a pattern of circular cutouts on both the holder and fixture align perfectly, and a vertical implant pin can be inserted through the cutouts to rigidly secure the fixture (Fig 4b). Furthermore, the holder’s simple cylindrical shaft makes it readily compatible with the mounting collets of most commercially available micromanipulators and easily adoptable on different surgical setups.

**Figure 4:**
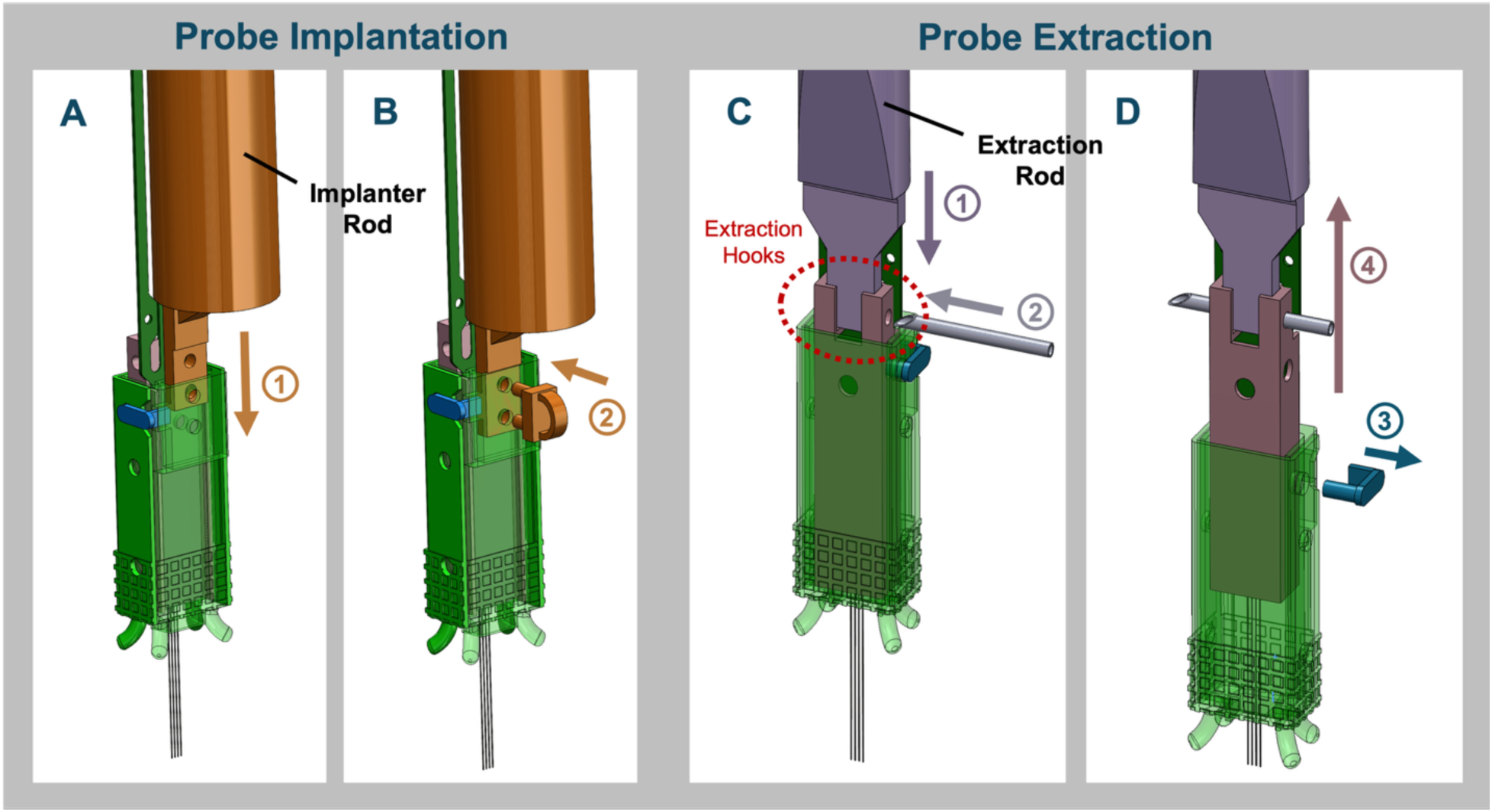
Implantation and Extraction. Implantation: (a)Inserting the tongue feature on the 3D printed implantation rod (orange) into the corresponding pocket on the back side of the bottom half outer casing (b)Once properly inserted, the two circular cutouts on both the implantation rod and outer casing lines up, allowing a small 3D-printed pin to slide in and secure the rod and hold it in place against the fixture assembly. Extraction: (c)Inserting the flat feature on the 3D printed extraction rod (purple) into the hooks of the inner sheath (pink). Once aligned, a 26-gauge hypodermic needle can thread through the inner sheath and extraction rod to lock them together (d)After removing the lock pin inserted during implantation (blue), retracting the extraction rod vertically will slowly pull up only the inner sheath-probe assembly from the rest of the fixture pieces that are adhered to the skull

After the probe has been implanted and the outer sheath is adhered to the skull, the holder is released by simply withdrawing the lock pin perpendicularly. This linear disengagement imparts minimal force on the implanted fixture along the most flexible axis of the shanks, ensuring the mechanical integrity of the probe is preserved and minimizing damage to the neural tissue of interest.

### Widespread Implementation and Success

To validate its robustness and usability, our fixture design was implemented across multiple laboratories, and here we demonstrate the stability results for four distinct experiments. These presented studies involved a total of 19 mice and 3 rats, with implant configurations ranging from one to four per animal, performed by researchers with varying degrees of surgical experience.

Across all experiments, the design demonstrated exceptional reliability, achieving a 100% success rate for implantation and a 100% retention rate for long-term functional recordings. Users consistently reported that the fixture assembly was easy to use and provided sufficient protection for the probe. Specifically, the two-part outer casing was highlighted as a key feature that significantly reduced the risk of accidental shank damage compared to designs that require the probe to be inserted shank-first. Furthermore, the screwless locking pin proved robust, providing sufficient rigidity to withstand insertion forces without failure, while allowing for a swift and gentle release of the probe holder post-implantation.

The stability of the fixture implants was confirmed in chronically implanted, freely-moving mice and rats. 93% of probes remained functional throughout the entire experimental timeline, ranging from 12 to 52 weeks, indicating the design securely buttresses the probe against forces exerted during naturalistic behaviors (e.g., locomotion, eating, and grooming). A detailed analysis of neural recording stability across time is presented in the following section.

### Compatibility with Diverse Experimental Hardware

A key advantage of our fixture is its compact, modular design, which ensures compatibility with the diverse and often crowded head-mounted hardware required for chronic experiments. Its minimal footprint allows it to be seamlessly integrated into complex assemblies alongside headplates, protective shrouds, and optical fibers without interference.

This versatility has been validated with the successful implantation of up to four probes simultaneously in a single freely moving mouse – a density and weight difficult to achieve with bulkier, heavier designs that may impair normal animal behavior. Furthermore, collaborating labs easily incorporated the fixture into their existing experimental setups, requiring no modification to their pre-designed skull plates or protective hardware. While all experiments in this paper were chronic implantations, the behavioral paradigm ranges from freely roaming exploration to head-fixed reaching tasks^14,15^, showcasing the wide range of experimental possibilities compatible with our fixture design.

These close-packed multi-probe trajectories are facilitated by pre-surgical CAD planning (Fig 5). We create a detailed 3D model of the skull to map precise trajectories for each probe, as well as the coordinates for the craniotomies and grounding screws. The fixture’s compact geometry makes it an optimal component for this virtual planning, enabling the design and precise execution of complex multi-probe arrangements.

**Figure 5:**
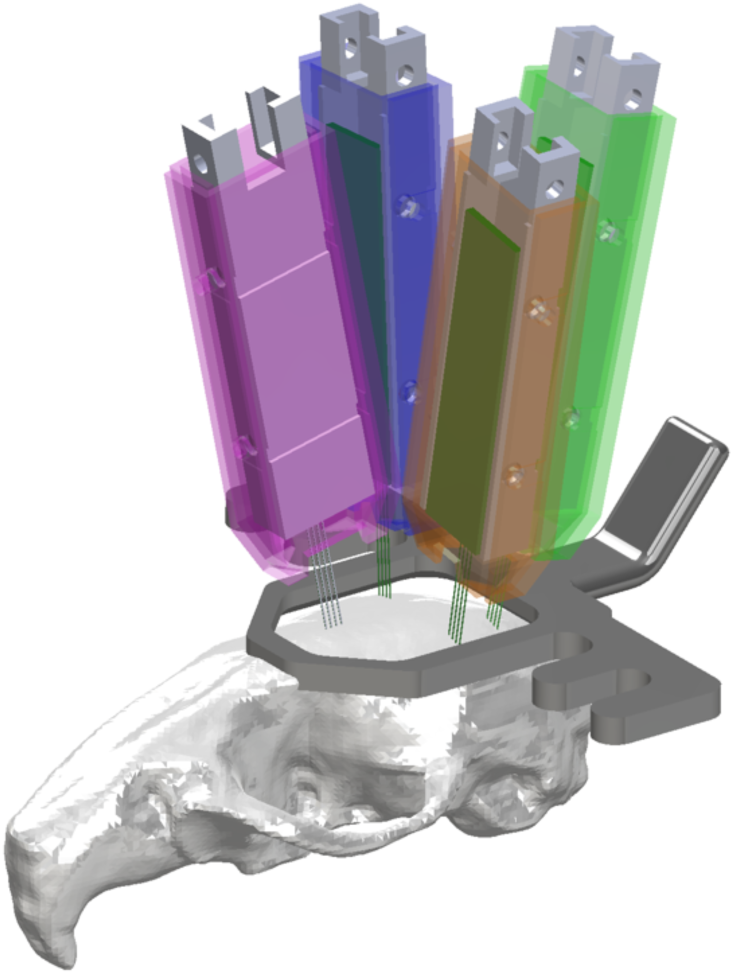
An example of CAD modeling of implantation trajectories on a mice skull including the physical models of our SCREWx fixtures. The modeling demonstrated here are the exact trajectories used in the 4-probe mice experiment in this paper. (Harris lab, surgeon A)

### Month-Long Stability of Chronic Recordings

To validate the long-term stability of recordings obtained with our fixture, we analyzed neural signals through (1) thresholded activity across all recording sites, and (2) yield of high-quality single units after spike sorting and curation. The thresholded event rate provides a baseline measure of signal integrity, independent of the variability introduced by spike-sorting algorithms or user-defined unit selection criteria. Our spike-sorting pipeline used Kilosort 4^16^ and accepted units that met standard, stringent quality metrics post curation^17^ (firing rate > 0.1 Hz, signal-to-noise ratio > 2, refractory period violations < 0.1, noise cutoff < 5, and presence ratio > 0.9)^18^. Both methods demonstrate that our fixture enables highly stable neural recordings over extended timescales, with experiments lasting up to a year without notable degradation in signal quality.

These stable recordings were achieved in 22 chronically implanted rodents (19 mice, 3 rats) across a wide range of brain regions, experimental paradigms, and labs, demonstrating the fixture’s versatility and ease of adoption. Freely-moving mice with single- and dual-probe implants were recorded during free exploration in an open field, social interaction tasks, Y-maze, and on an elevated plus maze, as well as during head-fixation with visual stimulation (Harris Lab). In head-fixed mice, recordings were obtained from subcortical brainstem neurons, specifically the vestibular nuclei, during controlled whole-body rotation experiments while the animals were passively rotated through space about the yaw axis to elicit neural responses (Cullen lab), providing a stringent test of implant rigidity and probe–brain coupling given the strong sensitivity of these neurons to head motion and mechanically induced perturbations. Mice with four-probe implants were also recorded during open field test, Y-maze, and novel object recognition tasks (Harris Lab). Rats were recorded during unrestrained behavior, including quiescent resting in the home cage and locomotion in a virtual reality dome. (Knierim Lab).

Qualitative inspection of the daily shank activity maps reveals a remarkably consistent pattern of high-amplitude activity, indicating stable proximity of the recording sites to active neurons (Fig 6 a-f). Aside from minor vertical drift, which is expected due to the small relative movements between the probe and brain tissue, the overall pattern of activity across shanks shows little degradation. These results are quantitatively supported by the stable yield of high-quality putative single neurons isolated through our spike-sorting and curation pipeline We sampled six recordings per animal from each experiment, spanning surgeries performed by different surgeons and performed the Mann-Kendall test to delineate whether there is a clear trend in the unit yield across time. The recordings used in this analysis were chosen to span roughly two weeks within the first 1.5 months since implantation, during which the animals are not undergoing any cognitive training or task. We found no statistically significant trend in any of the probe unit yield in these experiments. (all p-values > 0.05, results across 8 probes) (Fig 6g). Crucially, this consistency was observed across all animals and implant configurations, regardless of the probe’s insertion angle or target brain region. These results demonstrate that our fixture provides a secure and stable platform for a wide range of chronic recording experiments in both mice and rats. (See table 1 for full details)

**Figure 6:**
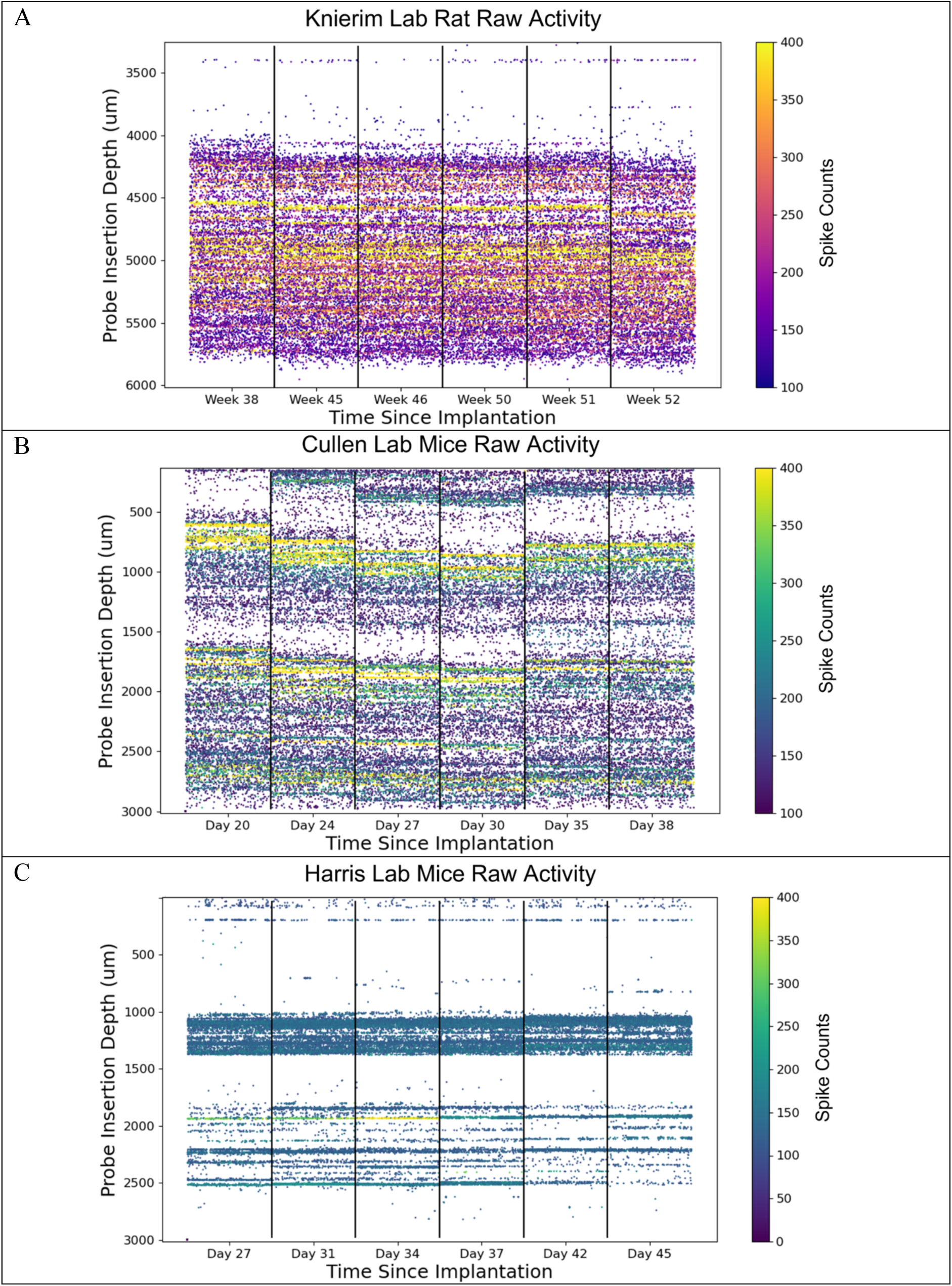
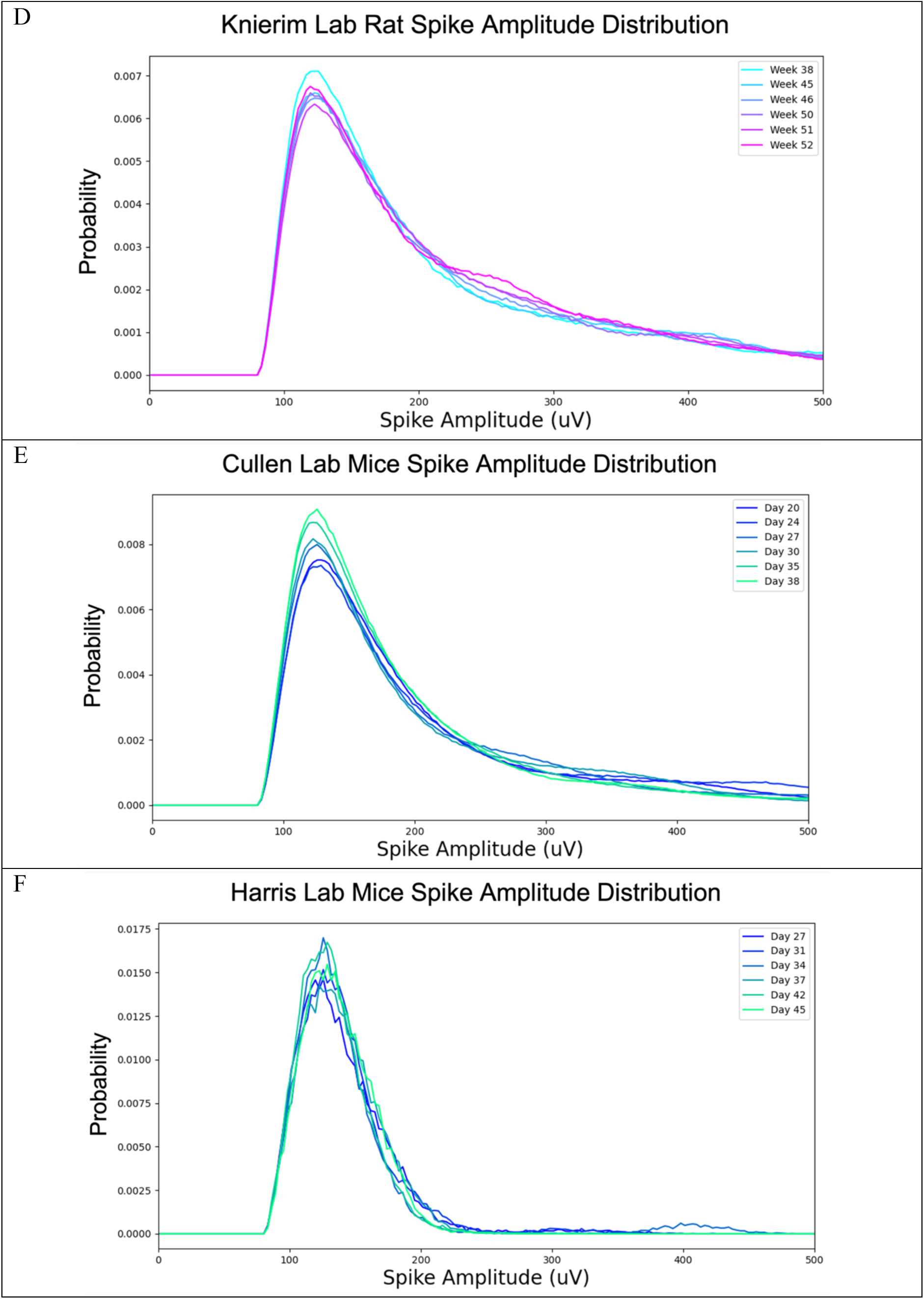
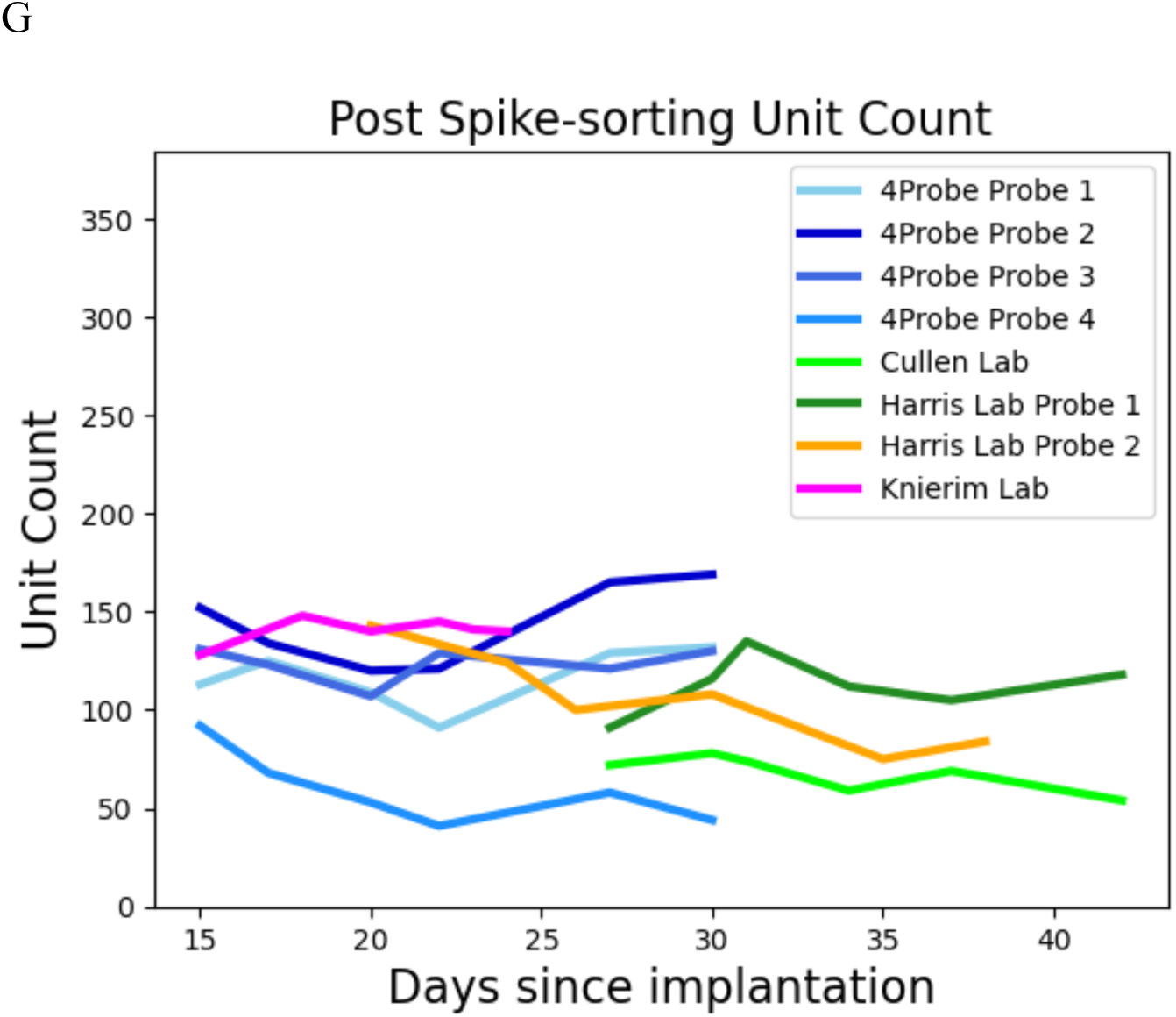
Stability of recordings across time. (a-c) Each panel shows a raw raster map of threshold-based spiking events across days from three different labs/ surgeons. Each column represents activity on a single shank with the most active sites during a 10min recording on a different day. The different time points reported here is due to the varying experimental timeline of different labs, with the longest one (Knierim lab) showing recording data one year after implantation(6a). (d-f) The corresponding amplitude distribution of the spiking events across days reported in the previous raster maps in figure 6a-6c. (g) Unit yield across days after spike-sorting and curation for the three animals reported in the previous figures, with the addition of a 4-probe mice from the Harris lab. Unit counts are reported independently for each probe with color groupings indicating to which animal they belong. No significant trend in unit count change across time was present in any of the recordings according to the Mann-Kendall test for each probe.

**Table 1:** Unit yield stability across days.

| Lab | Animal | Probe No. | Target Regions | Kendall's Tau | p-value |
| --- | --- | --- | --- | --- | --- |
| Knierim | Rat | 1 | RSC | -0.467 | 0.260 |
| Cullen | Mice | 1 | VN, PrP/NPH | -0.733 | 0.060 |
| Harris (2P) | Mice | 1 | mPFC | -0.600 | 0.133 |
| Harris (2P) | Mice | 2 | vHPC | 0.200 | 0.7071 |
| Harris (4P) | Mice | 1 | mPFC | 0.333 | 0.452 |
| Harris (4P) | Mice | 2 | RSA, dHPC, thalamus | 0.333 | 0.452 |
| Harris (4P) | Mice | 3 | S1, striatum | -0.067 | 1.000 |
| Harris (4P) | Mice | 4 | V1, vHPC | -0.600 | 0.133 |

### Minimal Impact on Naturalistic Freely-Moving Behavior

A critical requirement for any chronic implant is that it does not hinder naturalistic behavior. To confirm that our fixture’s low weight and compact dimensions met this standard, we compared locomotor activity in an open field arena in mice, as their low body weights make them more sensitive to the burden of head-mounted hardware, between one unimplanted animal and another two mice with a single and two chronically implanted probe(s) respectively. All three mice were of similar age, weight, and previous exposure to the open field apparatus. As demonstrated in Figure 7a, there were no qualitative differences in the locomotion pattern between the implanted and control group, as all started exploring the edge of the open field box before moving across the center of the maze. To confirm the added weight on the animals does not deter normal movement, we sampled their velocities within the first 600 seconds roaming freely in the open field maze and performed pairwise Wilcoxon rank-sum tests and reported no significant difference among the groups. (Fig 7b, all p-values > 0.05, results across three animals)

**Figure 7:**
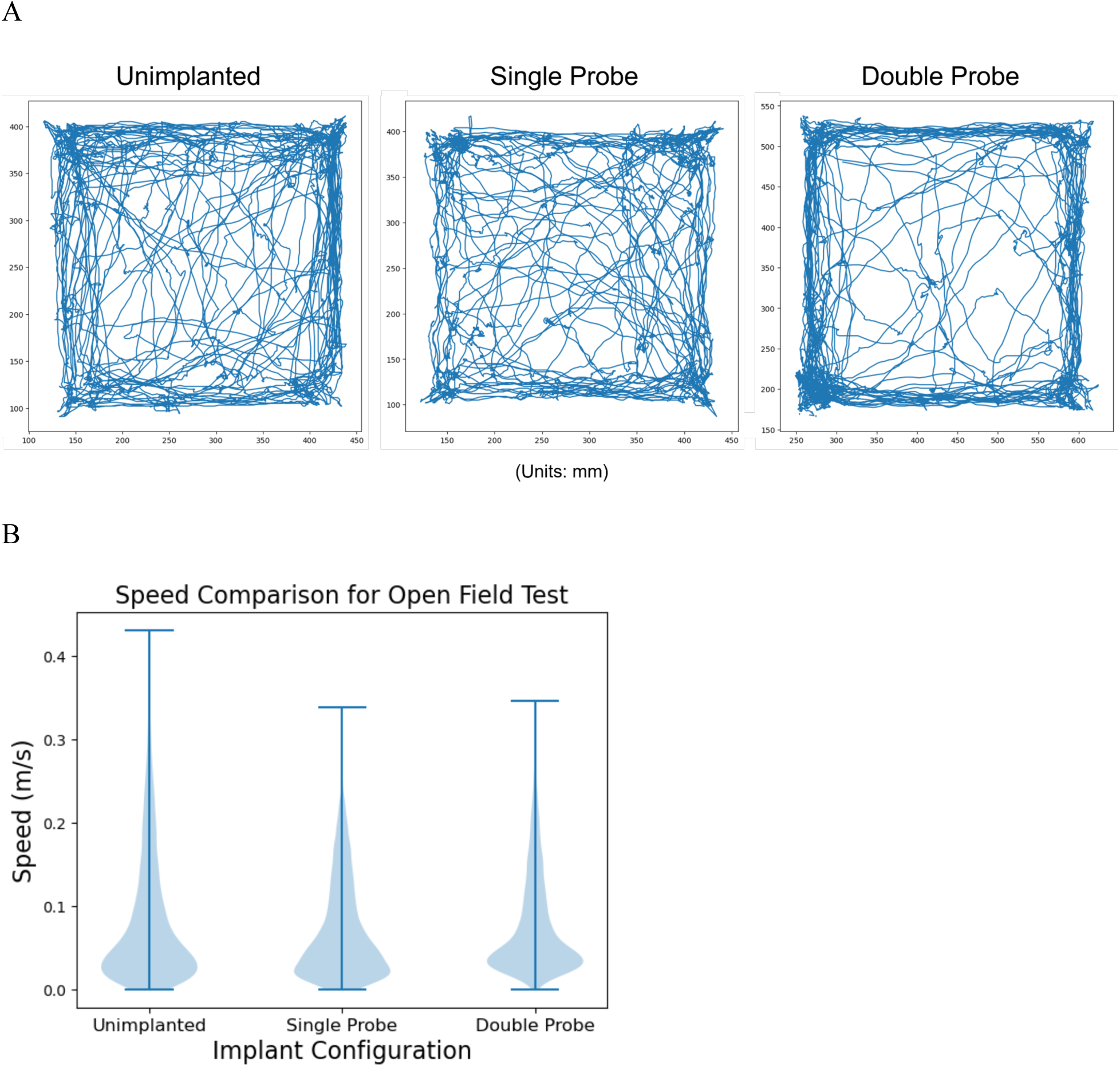
Locomotion in an open field test. (a) Trajectories of three mice in an open field test with no probes, 1 probe, and 2 probes respectively. Despite individual variance, the three mice exhibited highly stereotyped patterns of motion: surveying the edges upon first introduction to the apparatus and starting to move across the center of the open field as they spent more time familiarizing with the environment (b) Plot of velocities sampled across the first 600s in the open field test for the three mice. No statistical difference of median velocity was reported among the groups according to the pairwise Wilcoxon test.

This result indicates that the fixture is sufficiently lightweight and unobtrusive to permit unencumbered movement in small rodents. In addition to actively behaving during tasks, we monitored the implanted mice daily and observed they also exhibit normal feeding and drinking cadence, grooming, positive response to rodent enrichment, and frequent social interaction with their cage mates, all of which are indicative that the animals acclimate to added weight on their heads and resume normal behavior shortly after probe implantation. These data demonstrate this design is well-suited for chronic recording experiments that involve complex, naturalistic behaviors.

### Dedicated Probe Extraction and Recovery

A significant shortfall of many existing fixture designs is the lack of a dedicated, reproducible extraction mechanism. Probe recovery often relies on destructive or imprecise methods, such as drilling away cured adhesive or manually reversing the implantation process, both of which apply uncontrolled and potentially damaging forces to the probe. The design we report here overcomes these challenges by incorporating dedicated extraction features to enable controlled, standardized, and safe probe recovery. Two recovery hooks, integrated into the top of the inner sheath, remain exposed and accessible above the cemented outer casing throughout the experiment. To recover the probe, a specialized 3D-printed extraction tool is attached to a micromanipulator and carefully driven down to be seated between these hooks. The extraction tool is designed to precisely fit with the fixture hooks, restricting the motion between the inner sheath and the extraction tool. Aligned circular cutouts on the fixture and extraction rod allow a standard hypodermic needle to be inserted through these aligned holes, acting as a temporary shear-resistant tack to rigidly couple the inner sheath to the extraction tool (Fig 4c).

After the fixture’s lock pin is removed, the inner sheath and probe are mechanically decoupled from the outer casing. The entire assembly, now rigidly attached to the extraction tool, can be withdrawn slowly and precisely along the original insertion axis using the micromanipulator without off-axis deflections (Fig 4d). Overall, this controlled guided process minimizes unwanted forces, maximizing the likelihood of successful probe recovery.

### High-Yield Probe Recovery and Reuse

The ability to reuse costly Neuropixels probes makes resource-intensive, large-cohort size, and multi-probe experiments more feasible. To validate the reliability of our recovery method, we compiled extraction data from 30 probes across three surgeons with varying levels of experience.

We achieved an overall successful extraction rate of 83%. This high success rate was consistent across experienced operators (Surgeon T: 86%, n=21; Surgeon A: 100%, n=4; Surgeon M: 60%, n=5) and was not associated with systematic differences in fixture performance. The failures observed were not attributed to the fixture’s mechanical design. For instance, two failures (Surgeon M) were caused by pre-existing probe damage from hardened brain fluids caused by ground screw infections, a factor independent of the extraction process. The primary failure mode was breakage of a single shank at the brain’s surface during withdrawal. We hypothesize that this damage is not caused by the fixture itself, but rather by hardening of the sealant (e.g., Dura-Gel or Kwik-Sil) used to cover the craniotomy, which may adhere too strongly to the shank and cause it to break under tension during extraction.

Notably, the design proved effective for complex multi-probe recovery, as one surgeon successfully extracted all four probes from a single mouse after a 74-day chronic implant. Probes implanted in rats for long-term experiments of 12+ months have yet to be extracted.

Our criterion for a successful probe extraction and acceptable for reuse is based on the built-in probe BIST test in the SpikeGLX recording software. For Neuropixels 2.0 probes, this tests for the functionality of all the probe electronics, data transfer routes, as well as shank noise levels. A successful result in all of the categories tested indicates an intact probe that is still operating within the specifications given by the manufacturer. This confirms that implantation, long-term chronic recording, and extraction processes did not compromise the probe’s overall performance, confirming its eligibility for reuse.

## Discussion

Systems neuroscientists are increasingly interested in understanding how brain regions work together to orchestrate complex, naturalistic behaviors. Studying these questions often requires chronic, multi-probe recordings in freely-moving animals. To enable the feasible execution of such experiments, a need exists for fixture designs that enable stable long-term, multi-probe recordings and reliable probe recovery and reuse. Our novel fixture directly addresses these needs by overcoming critical limitations of weight, size, and risky handling procedures that have constrained previous designs.

Our design introduces four key innovations, all of which are improvements from the current state of the art: (1) small size/weight, (2) ease of assembly, (3) a screwless locking mechanism, and (4) dedicated extraction capability. First, at 0.4g and a cross section of 6.3 x 4.4 mm, the fixture is significantly lighter and more compact than most existing alternatives. These features enable high probe counts and proximity of neighboring probes to simultaneously target closely situated neuronal regions. As demonstrated by our successful four-probe implants in mice, the minimal footprint and light weight enable a density of implants previously difficult to achieve.

Second, our two-part encasing mechanism enables easy loading of fixtures without the need to glide the delicate probe shanks through a small opening, thus minimizing possible damage to probes.

Third, our screwless locking mechanism fundamentally improves the safety of probe implantation and release. Whereas incorporating screws produces a potential for torsional stress that poses significant risks of shank breakage, our design uses a locking pin that can be removed using only lateral force along a plane where the shanks are far more flexible and robust. Eliminating these off-axis forces inherent to previously published probe holders produced a more reliable fixture, which contributed directly to our 100% successful implantation rate and was frequently cited by collaborating users as a major usability improvement.

Finally, our fixture is the first to incorporate a dedicated extraction mechanism that is independent of implantation hardware. This mechanism allows for controlled probe recovery, crucial for achieving a high recovery success rate. This feature, combined with the screwless design, enabled our high 83% recovery rate, making routine reuse of probes practical.

The fixture’s performance was validated across multiple labs, animal models, brain regions, and experimental paradigms, demonstrating its versatility and robustness across a wide range of recording conditions. The design provided a mechanically stable platform for recordings lasting for at least three months and up to one year. Both the raw spiking activity and sorted single-unit yields showed remarkable consistency over time. This long-term stability demonstrates the fixture’s ability to support high-quality neural recordings in freely moving rodents for many months.

The high recovery rate and successful validation of reused probes in saline tests confirm that our system preserves the functional integrity of the probe. The ease of fixture fabrication using standard SLA 3D printers has already led to its adoption across multiple different groups^19^, demonstrating that it is a practical and accessible solution for the broader neuroscience community.

The primary failure mode for probe recovery was shank breakage at the brain’s surface, which we attribute not to a flaw in the fixture but to the procedural challenge of hardened brain fluids and dural sealants. Future work should focus on optimizing this aspect of the surgical procedure, perhaps by exploring alternative, less adhesive sealants or by developing micro-dissection techniques to dissolve or remove the sealant from the shank base before withdrawal without damaging underlying neural tissue.

In summary, we have developed and validated a lightweight, screwless chronic implant fixture that significantly lowers the barrier to entry for complex, multi-probe neuroscience experiments. By simplifying the handling process, minimizing implant weight, and dramatically improving the success rate of probe recovery, our design makes large-scale, longitudinal studies more accessible and cost-effective. Finally, we demonstrated that these fixtures can support high-quality, stable neural recordings from freely moving rodents for at least 3 months and up to one year. By sharing this tool freely and in an open-source manner, we hope to empower more researchers to investigate the multi-region neural dynamics that underlie complex behaviors.

## Acknowledgements

This work was supported by BRAIN initiative U01 NS115587 (T.D.H.) and National Institutes of Health (NIH) grant R01 NS039456 (J.J.K.).

## Notes

https://github.com/NP-HarrisLab/Harris-Lab-Neuropixels-Fixture-Design

### Competing Interest Statement

The authors have declared no competing interest.

### Summary of Updates

Added GitHub Links for design files and revised acknowledgements

